# Working Memory Capacities Neurally Dissociate: Evidence from Acute Stroke

**DOI:** 10.1101/2020.11.05.370486

**Authors:** Randi C Martin, Junhua Ding, A Cris Hamilton, Tatiana T Schnur

## Abstract

Substantial behavioral evidence implies the existence of separable working memory (WM) components for maintaining phonological and semantic information. In contrast, only a few studies have addressed the neural basis of phonological vs. semantic WM using functional neuroimaging and none has used a lesion-symptom mapping (LSM) approach. Here we address this gap, reporting a multivariate LSM study of phonological and semantic WM for 94 individuals at the acute stage of left hemisphere stroke. Testing at the acute stage avoids issues of brain reorganization and the adoption of patient strategies for task performance. The LSM analyses for each WM component controlled for the other WM component and semantic and phonological knowledge at the single word level. For phonological WM, the regions uncovered included the supramarginal gyrus, argued to be the site of phonological storage, and several cortical and subcortical regions plausibly related to inner rehearsal. For semantic WM, inferior frontal regions and the angular gyrus were uncovered. The findings thus provide converging evidence for separable systems for phonological and semantic WM that are distinguished from the systems supporting long-term knowledge representations in those domains.

## Introduction

Models of working memory (WM) in the verbal domain typically focus on the maintenance of phonological information (e.g. Gupta and Tisdale 2009; Page and Norris 1998). For instance, the well-known working memory model of Baddeley and colleagues (Baddeley et al. 2020) includes a phonological loop component, which consists of a buffer for maintaining phonological information and a rehearsal process that keeps this information active. However, considerable behavioral evidence from healthy and brain damaged individuals supports a multicomponent view of verbal WM, with separate buffers for maintaining phonological and semantic information (Martin et al. 1999; Shivde and Anderson 2011; see Martin et al., 2020, for review). Figure 1 shows a depiction of a model of WM delineating this approach. On the left-hand side are long-term knowledge representations for words, including their phonological and semantic information and, on the right, separate buffers for maintaining semantic and phonological information. Both for phonological and semantic information, long-term knowledge representations on the left are activated and stored in limited capacity WM buffers on the right^1^.

**Figure 1.**
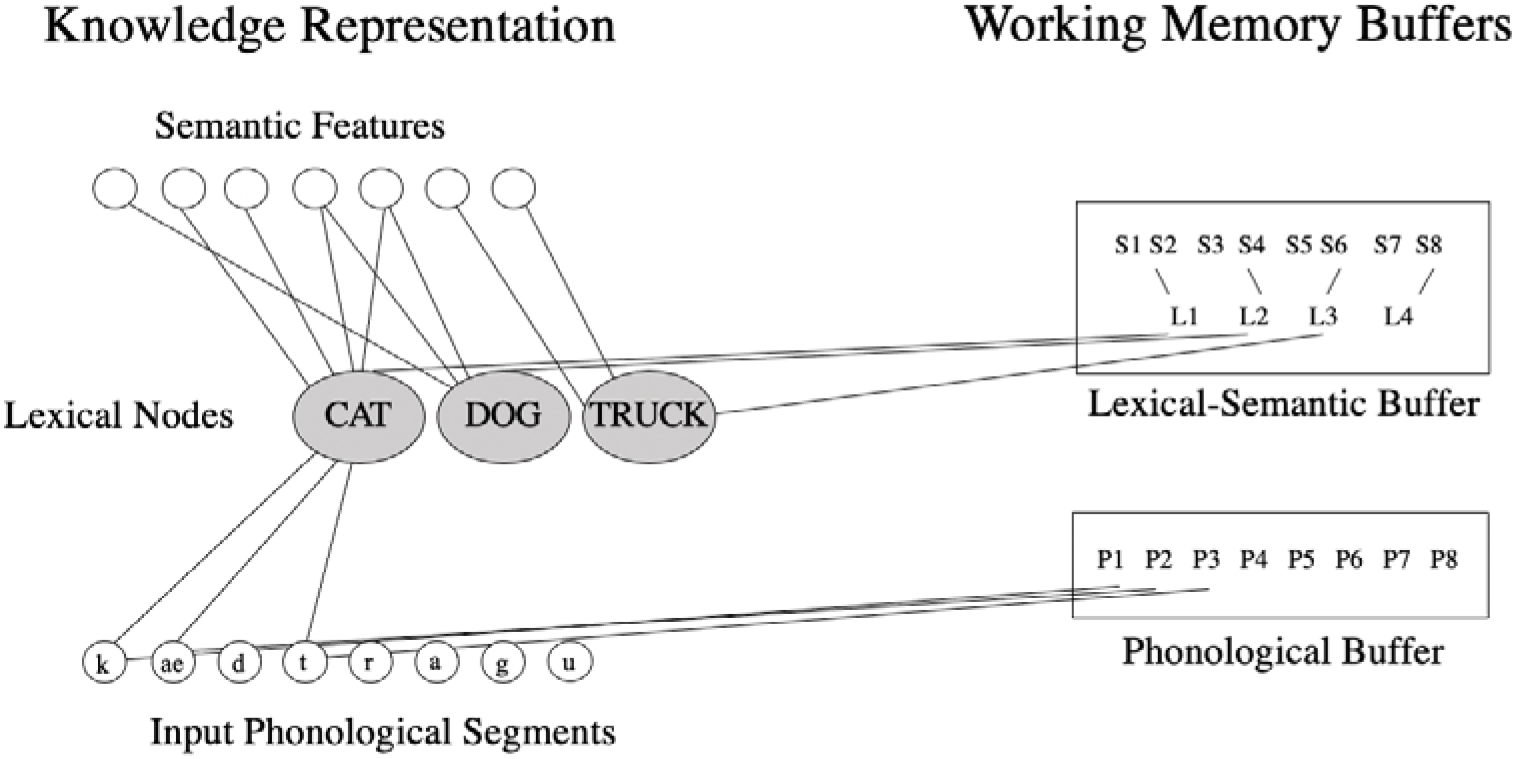
Model of working memory showing knowledge representations for words on the left and phonological and sematic working memory buffers on the right. (Based on Figure 1a from Martin et al. 2020.)

Consistent with the existence of separable WM capacities, the phonological and semantic components have been found to play different roles in sentence processing. The phonological component supports: 1) verbatim sentence repetition (Martin et al. 1994; Saffran and Marin 1975)^2^, and 2) speech rate in narrative production (Martin and Schnur 2019). In contrast, the semantic component supports: 1) the retention of word meanings prior to their integration across some distance during comprehension (e.g., maintaining the meaning of “cups” to integrate with the verb “cracked” in “Cups, vases, and mirrors cracked during the move”), and 2) the elaboration of phrases during sentence and narrative production (Martin and He 2004; Martin & Schnur, 2019; Potter 2012; Tan and Martin 2018; Tan et al. 2019). In contrast to a wealth of behavioral data suggesting separable capacities, limited evidence from functional or structural neuroimaging is available regarding differential neural bases of semantic and phonological WM capacities. The present study addresses this issue by using a multivariate lesion-symptom mapping (LSM) approach (Zhang et al. 2014; Mirman et al. 2015; Yourganov et al. 2016; Lacey et al. 2017; Schumacher et al. 2019), relating performance on tasks designed to tap semantic and phonological WM to the brain regions damaged in 94 individuals at the acute stage of left hemisphere (LH) stroke. Testing at the acute stage provides important advantages in lesion localization, as it avoids misinterpretation related to reorganization of function and the adoption of task strategies by the participants. By examining the relationship between lesion location and working memory performance, we demonstrate the brain regions required for phonological and semantic working memory, thus providing evidence as to the neural independence of working memory capacities. Below we summarize the current state of the literature regarding the brain basis of phonological and semantic WM.

### Left Inferior Parietal Localization of Phonological WM

As with behavioral studies, studies of the neural basis of verbal WM have focused on phonological WM, typically using tasks tapping immediate memory for random digit or letter lists. Early findings using lesion overlap or functional neuroimaging supported the conclusion that phonological WM was supported by an inferior parietal region (with the greatest overlap in the supramarginal gyrus; Paulesu et al. 1993; Shallice and Vallar 1990), which is separate from lateral superior temporal regions involved in the phonological processes underlying speech perception and word production (e.g., accessing the sounds associated with word meanings in order to understand or produce speech; Price 2012; Turkeltaub and Coslett 2010).

More recently, however, some authors have argued that phonological WM is inextricably linked to our long-term knowledge of phonology (N. Martin and Saffran 1997) and presented evidence that the same temporal lobe regions underlying phonological long-term knowledge support the temporary maintenance of phonological information (Ravizza et al. 2011; Leff et al. 2009). Such a view is consistent with embedded processes accounts of WM. Embedded processes accounts assume that, in general, WM consists of activated long-term knowledge representations together with a domain-general attentional system that brings some subset of activated information into the focus of attention (Cowan et al. 2020)^3^. In the phonological domain, various criticisms can be raised of these studies which claim a dependence of WM on long-term phonological knowledge. In neuropsychological studies some have claimed that the mechanisms that support phonological WM are the same as those that support phonological processes even for tasks with little or no WM demand (such as judging whether a stimulus is a word or nonword or accessing the meaning of a word from spoken input; Belleville et al. 2003; N. Martin and Saffran 1997). However, the tests of phonological processing often made some demand on phonological WM (e.g., syllable discrimination across a filled delay; N. Martin and Saffran 1997). Furthermore, Martin and Breedin (1992) demonstrated that individuals who were matched in having mild phonological processing deficits (e.g., scoring slightly below the control range in making word/nonword judgments to stimuli such as “pickle” and “bickle”) varied widely on phonological WM tasks requiring maintenance of digit lists (e.g., repeating back a list of 5 digits), from severely impaired to within the control range, supporting independence between processing and WM maintenance. In recent lesion overlap studies with large sample sizes, some studies have failed to control for patient speech perception abilities (Baldo et al. 2012; Ghaleh et al. 2020) or have factored out performance on tasks that arguably depend on phonological WM (Leff et al. 2009). Since phonological WM tests depend on speech perception, the degree of speech perception deficit should relate to patients’ WM performance. Further, phonological WM should correlate with the degree of damage to superior temporal regions based on the contribution of speech perception to performance on the WM task. The Leff et al. (2009) study controlled for nonword repetition which has been used as a measure of phonological WM capacity (e.g., Gathercole and Baddeley 1989), and thus its removal may have taken out a large portion of the variance due to phonological WM. Recently, in a large sample study of patients undergoing glioma resection, Pisoni et al. (2019) specifically contrasted phonological processing and WM regions and found only partial overlap between them, with damage to the supramarginal gyrus related only to WM. Thus, the neuropsychological evidence to date supporting a dependence of WM on long-term phonological knowledge is difficult to interpret given the methodological confounds.

Criticisms can also be raised with respect to fMRI studies claiming a reliance of phonological WM on phonological processing regions. One issue is that the speech perception regions have not typically been established in the same participants carrying out the phonological WM test, and sometimes the regions assumed to support phonological processing are remote from those indicated in meta-analyses of speech perception (e.g., Ravizza et al. 2011). Also, most studies have used visually rather than auditorily presented word lists (e.g., Ravizza et al. 2011; Lewis-Peacock et al. 2012). Although there is substantial evidence that visual verbal stimuli are recoded phonologically during WM tasks, one would expect more direct and consistent activation of phonological codes from auditory input. With visual input, participants may, at least on some trials, rely on memory for visual or orthographic features of the stimuli, reducing the sensitivity in detecting regions involved in phonological maintenance. Recently, Yue et al. (2018) carried out an fMRI study using a probe recognition task with auditorily presented lists of nonword stimuli (e.g., list: treb, plim, suke; probe: trem). They found that the SMG showed activation and WM load effects during a delay period between the list and the probe whereas the superior temporal region identified for the same individuals as supporting speech perception did not. Moreover, using multivariate decoding methods (MVPA), phonological information could be decoded in the SMG irrespective of the classifier used whereas in the STG, decoding was successful for only one classifier, and moreover, decoding accuracy across individuals in the SMG correlated with their WM performance whereas decoding accuracy in the STG did not. In sum, the differential evidence for similar or different regions involved in phonological long-term knowledge vs. its maintenance may be the result of the use of tasks which did not strongly require phonological working memory, did not control for phonological input processing, or the assessment of phonological long-term knowledge using tasks that also required working memory.

### Left Inferior Frontal Involvement in Semantic WM

In comparison to phonological WM, relatively little is known about the neural basis of semantic WM. Substantial evidence indicates that long-term semantic knowledge is represented in bilateral middle and inferior temporal lobes (e.g., Mummery et al. 1999; Visser et al. 2012). With regard to semantic WM, Martin (2005) noted that individuals identified as having semantic WM deficits had lesions encompassing left inferior frontal regions, which distinguished them from those with disruptions of semantic knowledge per se (Mummery et al. 1999) and from those with phonological WM deficits and inferior parietal lesions (Shallice and Vallar 1990). These findings would again argue against an embedded processes approach to WM, given the different localization of the WM and LTM regions for semantic WM. However, large sample studies of brain damaged individuals have not been carried out examining the regions involved in semantic WM maintenance, while controlling for semantic knowledge deficits.

In early fMRI studies with healthy subjects, Martin et al. (2003) and Shivde and Thompson-Schill (2004) contrasted performance on short-term memory probe tasks tapping WM for phonological (rhyme or vowel probe) vs semantic (synonym probe) information (e.g., subjects heard a list of 1 or 4 words and, after a short delay, answered whether a probe stimulus was similar in rhyme/vowel/or meaning with one of the list words). Both reported that left parietal regions were more activated for the phonological than the semantic task whereas inferior/middle frontal regions were more activated for the semantic tasks. Hamilton et al. (2009) examined regions involved in maintaining word meanings prior to their integration in a task contrasting high and low demands on semantic WM. In this study, subjects judged whether adjectives could be sensibly integrated with a noun (e.g., green emerald vs. green sun). In the high WM demand condition, adjectives came before a noun (e.g., green, shining, bright emerald/sun) whereas in the low demand condition, the adjectives came after the noun (e.g., emerald/sun bright shining, green). The logic was that in the “before” condition, the meanings of the adjectives had to be maintained until the noun was processed, whereas in the “after” condition, each adjective could be integrated with the noun as it was perceived. Again, left inferior/middle frontal regions were more activated in the high than the low semantic WM demand condition. In an fMRI study using MVPA, Lewis-Peacock et al. (2012; Exp 2) found that semantic maintenance could be discriminated from phonological and visual maintenance in left anterior frontal and superior temporal regions. (It should be noted though that their stimuli were presented visually and the regions showing the greatest differentiation of phonological from semantic and visual maintenance were in bilateral occipital lobes, suggesting that participants may have relied on orthographic coding to complete the supposedly phonological task.) In sum, there are only limited findings regarding the neural basis of semantic WM and those that exist tend to suggest the involvement of a left inferior frontal region.

### Complicating Factors: Additional Regions Involved in Phonological and Semantic WM

The above findings are consistent with the claim of a contrast between left frontal regions supporting semantic WM and left parietal regions supporting phonological WM. However, this claim is complicated by other findings indicating a role for left frontal regions in articulatory rehearsal and a role of a left parietal region (i.e., the angular gyrus) in semantic processes (Binder et al. 2009). With regard to rehearsal, a long-standing assumption has been that subvocal articulatory rehearsal is used to support the maintenance of phonological forms (Baddeley et al. 1975), and rehearsal is a major component of the phonological loop in Baddeley et al.’s (2020) model. In the Yue et al. (2018) neuroimaging study, left frontal regions (including the precentral gyrus, posterior inferior frontal gyrus [IFG], and supplementary motor area) showed a load effect during the delay period of the phonological WM task. Previous imaging studies have provided evidence that these regions are involved in either subvocal rehearsal or executive processes related to motor planning (Chein and Fiez 2001; Smith et al. 1998). In Yue et al. 2018, the putamen and cerebellum also showed load effects, and these regions are also likely involved in rehearsal given their role in controlling motor aspects of speech production. With regard to left parietal regions’ involvement in semantic maintenance, many neuroimaging and neuropsychological studies have reported evidence that the angular gyrus (AG) plays an important role in semantic processing (Bemis and Pylkkanen 2013; Binder et al. 2009; Jefferies 2013; Price et al. 2015), with some of that evidence pointing to a role in semantic WM, given its contribution in integrating word meanings during phrase or sentence comprehension (e.g., Humphreys et al. 2007; Price et al. 2015). Yue et al. (2018) found evidence supporting a role for the AG in semantic WM as he showed that semantic representations could be decoded in the AG during the delay period of a semantic WM task, which involved judging the relatedness of word meanings across a delay.

Thus, there is strong evidence suggesting both a frontal-parietal dissociation for phonological vs. semantic working memory as well as the reverse. Although the frontal areas proposed to be involved in semantic WM (left inferior/middle frontal gyrus) differ from those proposed to be involved in subvocal articulatory rehearsal (more posterior left inferior frontal, precentral gyrus, and supplementary motor areas; e.g., Smith et al. 1998), some of these areas overlap and are not clearly distinguished based on early fMRI results (e.g., LIFG involvement in rehearsal from Chein and Feiz 2001, and LIFG involvement in semantic WM from Shivde and Thompson-Schill 2004). Similarly, the left inferior supramarginal gyrus proposed to be involved in phonological WM is bordered by the left angular gyrus, which potentially plays a role in semantic WM and prior neuroimaging studies have not directly contrasted the roles of these two neighboring regions in semantic vs. phonological WM. Furthermore, because activation in a region revealed through functional neuroimaging approaches does not establish its necessity in processing (e.g. Rorden and Karnath 2004), it is important to have converging lesion-based data, where behavioral impairments following damage to a region strongly implicate its necessary role. Thus, the current study provides critical data through examining the neural basis of semantic and phonological WM in the same subjects using a multivariate LSM approach where we have lesion coverage for these regions to determine if they can truly be differentiated regarding the type of WM that they support.

### Current Study

To address the distinctiveness of the brain regions involved in phonological and semantic WM, we used multivariate LSM to relate disruption of phonological and semantic WM to brain damage in acute LH stroke, while controlling for individuals’ single word phonological and semantic processing abilities. To test phonological WM, we used a digit matching span task, in which participants heard two lists of digits and decided whether they matched (Martin et al. 1994; Tan and Martin 2018). Digit lists were used as there is relatively little semantic information conveyed by random sets of digits. To test semantic WM, we used a category probe task in which participants judged whether a probe word was in the same semantic category as any list word (Martin et al. 1994; Tan and Martin 2018). Neither task required speech output, thus avoiding contributions of overt speech production deficits to WM performance. Both of these measures have been used in prior behavioral studies and have been found to relate to different aspects of language comprehension and production, as discussed earlier (e.g., Martin and He 2004). While these measures tap into different capacities, prior results have shown a significant correlation between them (Tan and Martin 2018; Tan et al. 2017), which is unsurprising in that an ability to retain phonological information would help to support performance on the category probe task. That is, even if semantic representations of the to-be-remembered items had been lost by the time the probe was presented, a surviving phonological record could be used to re-access semantics. In the other direction, there is evidence for a boost from semantics, or at least familiarity, in digit span tasks, in that lists containing subsequences of digits that are more familiar (e.g., 1492) are better recalled (Jones and Macken 2015). Thus, to determine regions specific to one of the WM capacities we factored out performance on the other WM task in our analyses. In order to control for participants’ speech perception and semantic knowledge, we also factored out performance on a picture-word matching task with semantically and phonologically related distractors (Martin and Schnur, 2019). To our knowledge this is the first study to examine the neural dissociation between phonological and semantic WM in a large group of persons with acute LH stroke while accounting for previous confounds of phonological input processing, semantic knowledge, the covariation between measures of the two capacities, and reorganization of function.

## Materials and Methods

### Participants

Ninety-four acute left hemisphere stroke patients (51 males; 81 right-handed; 88 ischemic stroke; 6 hemorrhagic stroke; age: M = 61 years; S.D. = 12 years; range = 25-85 years; education: M = 14 years; S.D. = 4 years; range = 0-33 years) were recruited from the Memorial Hermann, Houston Methodist and St. Luke’s hospitals in Houston, Texas, USA, as part of an ongoing project (Martin and Schnur 2019; Ding et al. 2020). Subjects met the following inclusion criteria: Native English speaker; No other neurological diseases (e.g., tumor, dementia); No neuroradiological evidence of previous non-lacunar left hemisphere stroke (cf. Corbetta et al. 2015). Behavioral testing and clinical neuroimaging completed within one week after stroke (7 subjects within two weeks; median = 3 days, range = 1-12 days). We recruited 13 non-brain damaged participants as controls (three males; 11 right-handed; age: M = 55 years; S.D. = 14 years; range = 37-78 years; education: M = 16 years; S.D. = 3 years; range = 12-22 years) matched to patients on demographic variables (age: *t*(105) = −1.53, *p* = 0.13; education: *t*(86) = 1.66, *p* = 0.10; handedness: *x*^2^ = 0.02, *p* = 0.88). Informed consent was approved by the Baylor College of Medicine Institutional Review Board.

### Behavioral tests

#### Phonological working memory

We measured phonological working memory (WM) with the digit matching span task (Allen et al. 2012; Martin and Schnur 2019; four participants completed the digit span task using the standard WAIS-R procedure, Wechsler and De Lemos 1981; Allen et al. 2012; Martin and Schnur 2019). With respect to digit matching span, participants first heard two-digit lists, in which one digit was presented per second. Participants judged (yes or no) whether the two lists were the same or not. In the ‘non-match’ trials, the second list reversed one pair of two adjacent digits (e.g., 5 3 1 8 – 5 1 3 8). This reversed position was randomized. List length increased from 2-6 digits, with 6, 8, 6, 8, and 10 trials per list length, respectively. Half of the trials matched, half did not. Different numbers of lists were used per list length such that the position of the reversal was approximately equal across serial positions within each list length, while keeping the number of lists low overall. We stopped testing when participants’ accuracy fell below 75% for a particular list length. We calculated the phonological WM span dependent measure for each list length by linear interpolation between the accuracy of the two list lengths spanning 75% accuracy. For example, if the subject scored 7/8 (87.5%) correct at list length 3 and 4/6 (66.7%) correct at list length 4, then span would be 3 + ((87.5-75)/(87.5-66.7)) = 3,60. If accuracy for the two-digit lists was lower than 75%, we assumed 100% accuracy for a 1-item list length. If accuracy for the six-digit lists was higher than 75%, we assumed 50% accuracy for a 7-item list length.

#### Semantic working memory

We measured semantic WM using the category probe task (Martin et al. 1994; Martin and Schnur 2019). Participants judged whether a spoken probe word was in the same semantic category as any word in a preceding spoken list. The categories were animals, body parts, clothing, fruit and kitchen equipment. For example, for the list: bear dress apple leg and probe: pear, the answer would be “yes,” since pear and apple are in the same category (fruit). The matched position in lists was randomized. List length increased from 1 to 4 items, with 8, 8, 12 and 16 trials per list length, respectively. On half of the trials the probe matched the category of one of the list items whereas on the other half it did not. The number of trials per list length varied such that the position of the matching word was approximately equal across serial position within each list length, while keeping the number of trials low overall. We presented words one per second. We stopped task administration when participants’ accuracy was below 75% for a given list length. Scoring proceeded as for digit matching span.

#### Phonological and semantic input processing

We measured the integrity of phonological and semantic input processing via a word picture matching task (Martin et al. 1999; Breese and Hillis 2004; Martin and Schnur 2019). We presented seventeen pictures four times. Each picture was presented with an auditorily presented matching word, phonologically related foil, semantically related foil and an unrelated foil, for a total of 68 trials distributed across four different presentation sets. Participants judged (yes/no, verbally or non-verbally) whether the picture and word represented the same object. We calculated phonological and semantic d’ scores to estimate participants’ ability to discriminate between matching trials, and phonological and semantic foil trials. We used a composite measure of input processing because the phonological and semantic input processing scores were highly correlated (*r* = 0.69; *p* < 0.001). The composite measure was generated by a principal component analysis including phonological and semantic input processing d’ scores (explained variance = 85%; phonological input processing loading = 0.92; semantic input processing loading = 0.92).

### Image acquisition and preprocessing

We identified participants’ lesions from diffusion weighted, T1, T2 FLAIR images (scanned in the axial direction) and for those contraindicated for MRI, CT scans. Neuroimaging was collected within 1.5 days of behavioral testing (range 0-8 days). The resolution of diffusion-weighted and T1/T2 images was 1*1*4.5 mm and 0.5*0.5*4.5 mm, respectively and 0.5*0.5*5.0 mm for CT scans (n = 6 subjects).

To quantify patients’ lesions, we first registered diffusion weighted images to T1/T2 images using AFNI (https://afni.nimh.nih.gov/). Then lesions were demarcated on the diffusion weighted images using ITK-snap (https://www.itksnap.org/pmwiki/pmwiki.php). Finally, we normalized T1/T2 and lesion images into the MNI space using ANTS (https://stnava.github.io/ANTs/; Avants et al. 2008). With regard to CT images, lesions were demarcated directly on the Colin-27 template while referring to CT images.

### Multivariate LSM

To identify lesion location patterns associated with phonological and semantic WM, we conducted support-vector regression (SVR; libsvm 3; https://www.csie.ntu.edu.tw/~cjlin/libsvm/) LSM using MATLAB 2018b (https://www.mathworks.com/products/matlab.html). To control for potential confounding factors, we measured the relationship between brain damage location and either phonological or semantic WM performance independent of the contribution to performance from lesion size (cf. Sperber and Karnath 2018; DeMarco and Turkeltaub 2018), input processing deficits, and the opposing working memory ability. Specifically, we calculated the residuals of phonological and semantic WM scores by regressing out lesion size (number of lesion mask voxels), the composite measure of input processing and the other WM score. We controlled for the other WM score because the two WM measures were significantly correlated (*r* = 0.51; *p* < 0.001) and we wished to determine the relation to brain areas for the component of these measures specific to either semantics or phonology. We did not control for demographic variables (age, education, sex, handedness and days tested post-stroke) because correlations with the two WM scores were not significant after multiple-comparison correction (Bonferroni corrected *p* > 0.27). In order to normalize dependent variables (Zhang et al. 2014), the residuals were further scaled ((value-min)/(max-min)) to fall within the continuous range of 0-1 (Hsu et al. 2003). We only included voxels lesioned in at least 5 (5%) of 94 patients. Voxels with the same lesion pattern across patients were combined as a patch (Pustina et al. 2018). We used all the patches to predict working memory residuals. A grid search was conducted to select the optimal parameters (cost: 0.01 – 10^9 and gamma: 10^-9-1000; the same range as scikit-leran: https://scikit-learn.org/stable/index.html) for rbf-kernel SVR (Mirman et al. 2015). For each parameter combination, we used 5-fold cross-validation to acquire its mean square error of real and predicted dependent scores. We used 1000 permutations (shuffling dependent scores) to acquire the parameter combination significance level (the rank of the real model’s mean square error in 1000 random models). The parameter pair with the lowest *p* value was considered the optimal combination which we then used in subsequent analysis. Finally, patterns of brain region damage significantly related to WM performance were determined via permutation test (1000 iterations) using the optimal parameter pair model (Zhang et al. 2014). The significance level of each patch was the rank of its beta value in beta values of 1000 random models. Only negative beta values were of interest (i.e., our expected direction). We set the significance threshold to *p* < 0.05 for models and patches. We defined brain regions based on the brain connectome atlas (Fan et al. 2016). The brain map was generated by REST (http://www.restfmri.net/forum/; Song et al. 2011). To note, consistent with previous literature and the brain connectome atlas’s gyral subdivisions, we subdivided the insula using an anatomical solution (i.e. anterior dorsal, anterior ventral and posterior insula; Deen et al. 2010) as the brain connectome atlas’s division of the insula uses cytoarchitectonic nomenclature.

## Results

### Behavior

Compared with the non-brain damaged participants, patients showed significant impairments on phonological WM (span length scores; controls: M = 6.12, S.D. = 0.50, range = 5.00-6.38; patients: M = 4.94, S.D. = 1.59, range = 1.37-6.50; *t*(55) = 5.48, *p* < 0.0001), semantic WM (span length scores; controls: M = 4.59, S.D. = 0.80, range = 3.35-5.45; patients: M = 2.16, S.D. = 1.38, range = 0.33-4.50; *t*(23) = 9.22, *p* < 0.0001), phonological input processing (d’ scores; controls: M = 3.70, S.D. = 0.19, range = 3.44-3.78; patients: M = 3.14, S.D. = 0.71, range = −0.15-3.78; *t*(92) = 6.69, *p* < 0.0001) and semantic input processing (d’ scores; controls: M = 3.18, S.D. = 0.39, range = 2.60-3.78; patients: M = 2.70, S.D. = 0.74, range = 0-3.78; *t*(105) =2.32, *p* = 0.02; see Figure2).

**Figure 2.**
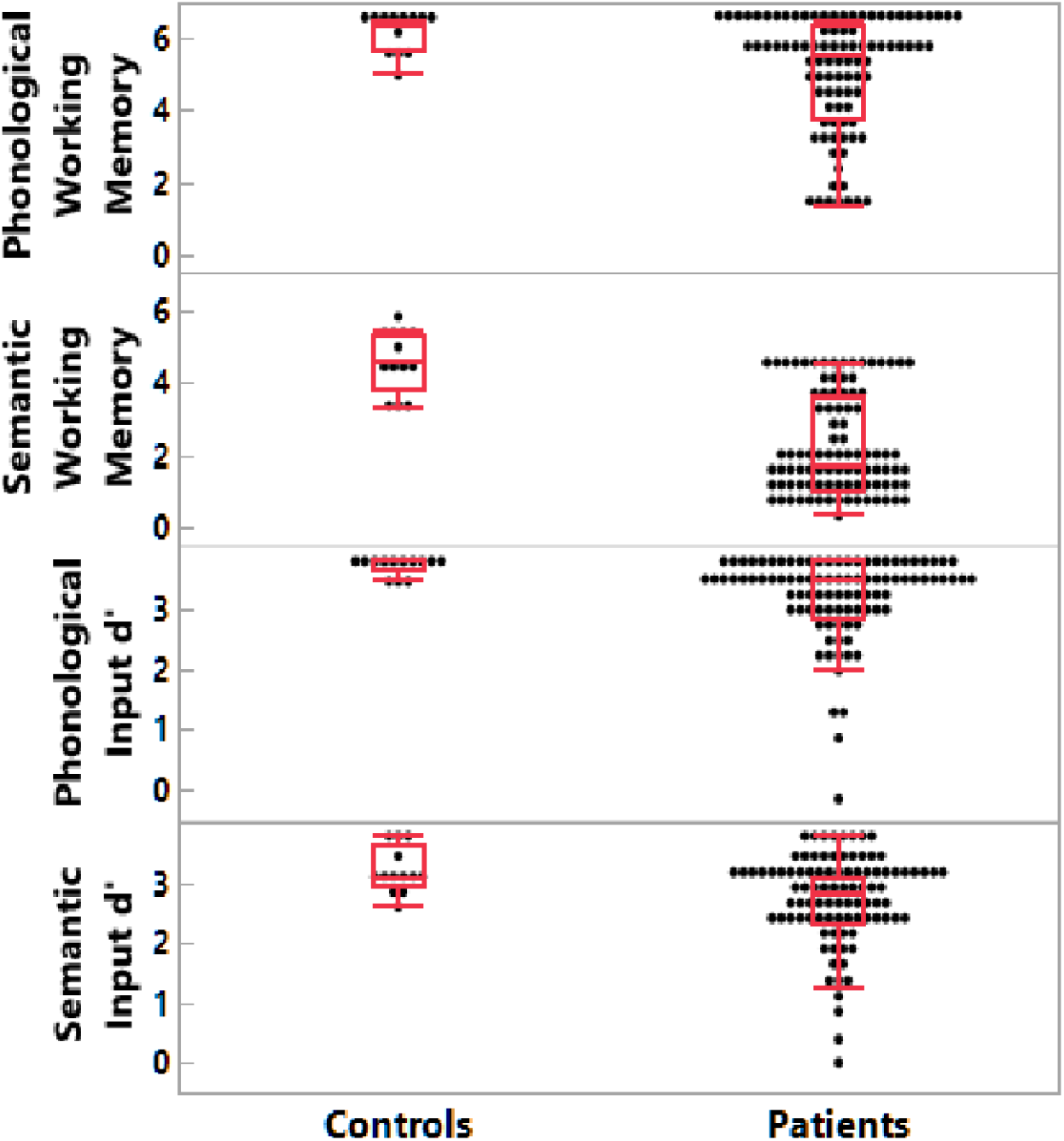
Patient and non-brain damaged control behavioral results. A) Phonological working memory span; B) Semantic working memory span; C) Phonological input processing; D) Semantic input processing.

### Lesion distribution

Figure 3A displays lesion coverage across patients (lesion size: M = 14098 mm^3^, S.D. = 18641 mm^3^, range = 135-104243 mm^3^). The primary damaged regions (regions damaged in at least five subjects and with > 100 lesioned voxels) included the middle and inferior frontal gyri, pre- and post-central gyri, superior temporal gyrus and sulcus, superior and inferior parietal lobules, insula, basal ganglia and thalamus. Figure 4 shows correlations between regional proportion damage and the distances between the damaged regions. Figure 4 reveals lower correlations in proportion damage between remote regions, indicating feasibility to test for functional dissociations between remote regions (e.g. SMG/ BA 39 vs IFG/ BA 44, 45). Moreover, not all adjacent damaged regions presented with higher correlations between proportion damage, indicating feasibility to test for functional dissociations between adjacent regions’ functions (e.g. SMG/ BA 39 vs AG/ BA 40).

**Figure 3.**
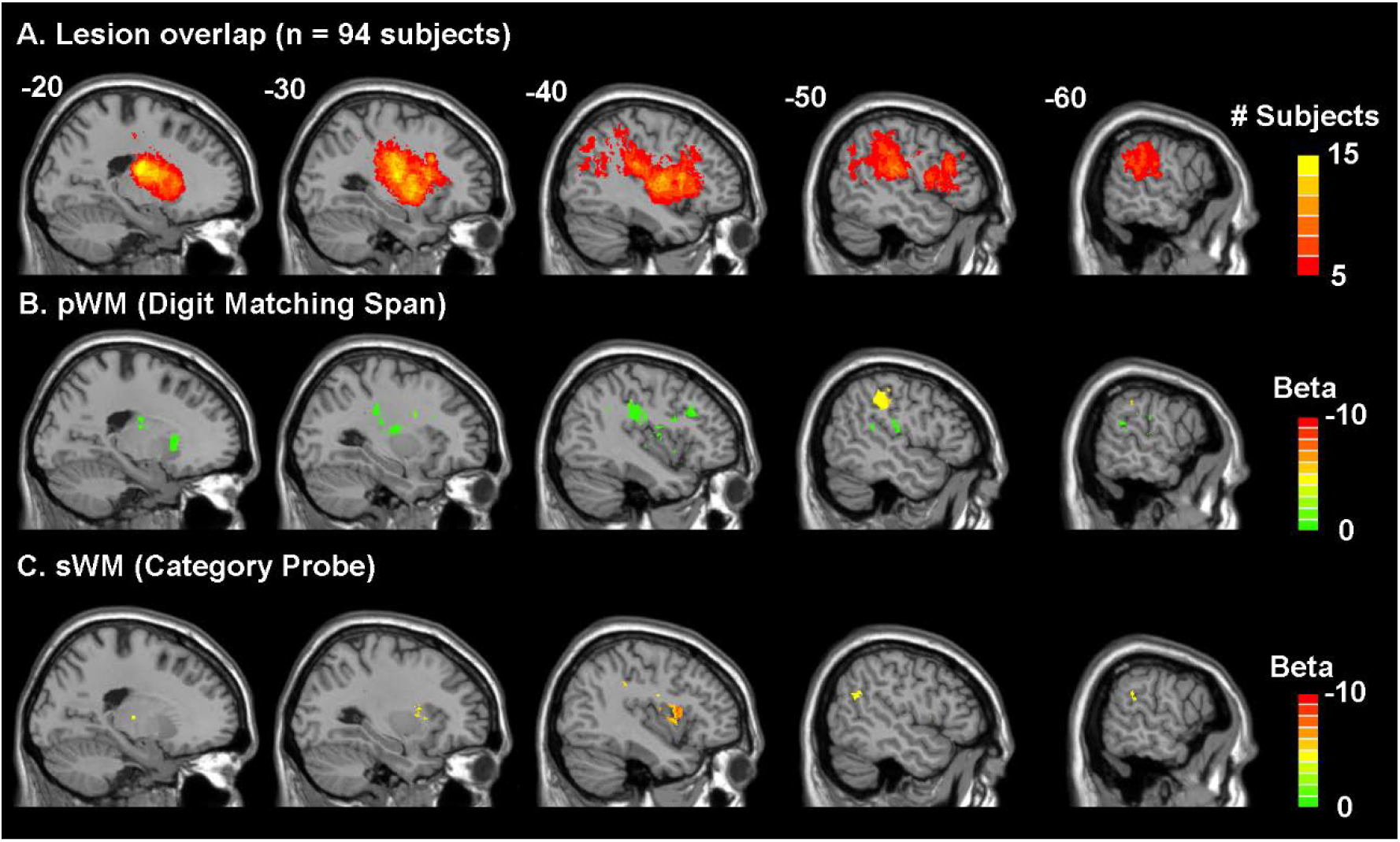
Lesion Overlap Distribution (A) and Lesion-Symptom Mapping Results (B & C). A) Lesion overlap in 94 acute left hemisphere stroke subjects where only regions damaged in at least five subjects (> 5%) were included in the lesion-symptom mapping analyses. Figure 3B and Figure 3C show the beta values of the regions significantly associated with decreased performance in the phonological WM (Figure 3B) and semantic WM (Figure 3C) measures after accounting for lesion volume, input processing (input processing composite score of semantically- and phonologically related word-picture matching d’ scores), and the respective opposing working memory task (*p*’s < 0.05). p/sWM: phonological/semantic working memory.

**Figure 4.**
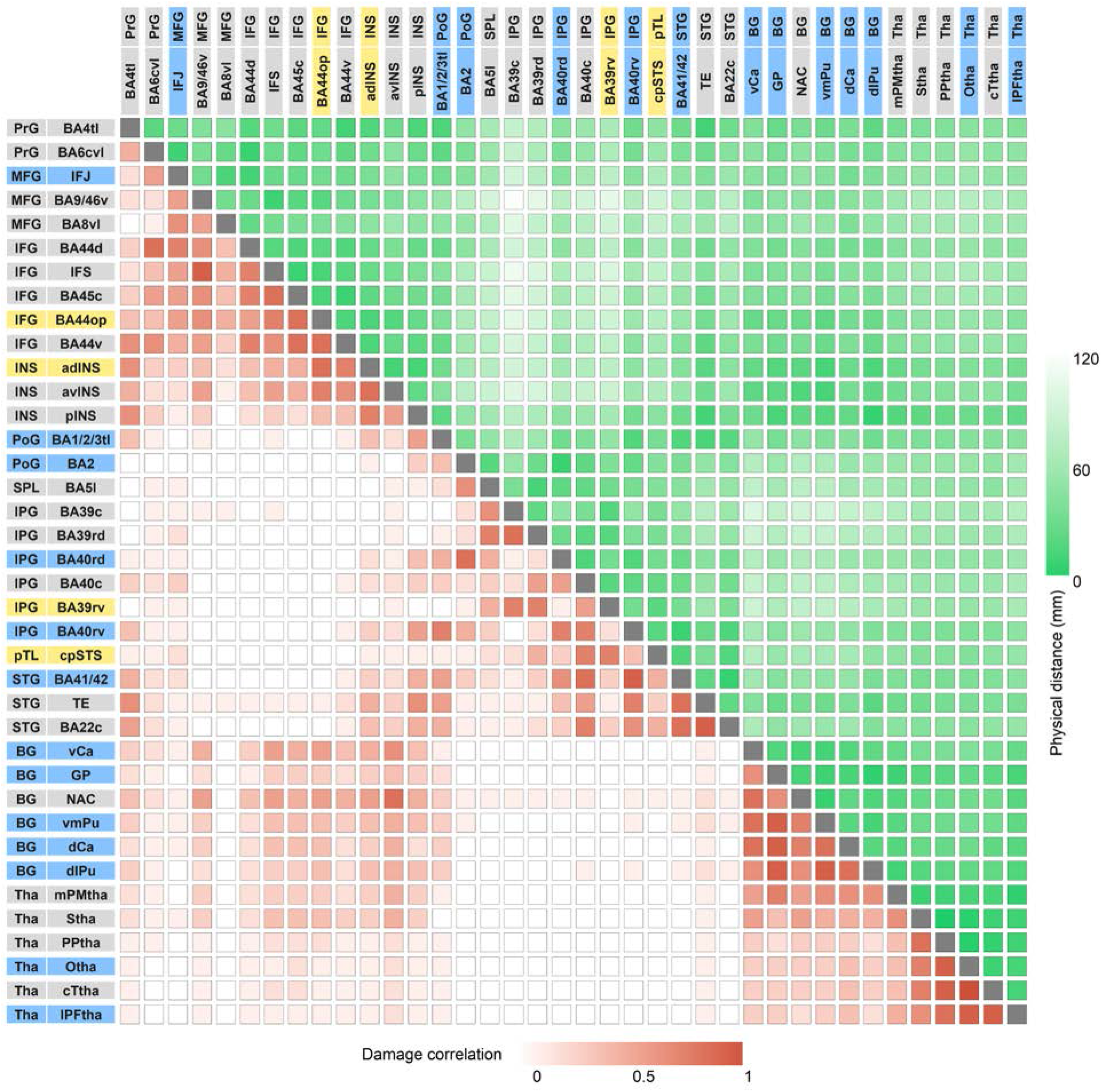
Matrix of between region damage correlations and distances. Increasing color intensity reflects either increasing correlations of proportion damage between regions (in red) or decreasing distance between regions (in green). Blue and yellow labels indicate the significant regions from the phonological and semantic WM LSM analyses, respectively. PrG: precentral gyrus; MFG: middle frontal gyrus; IFG: inferior frontal gyrus; INS: insula; PoG: postcentral gyrus; SPL: superior parietal lobe; IPL: inferior parietal lobe; pTL: posterior temporal lobe; STG: superior temporal gyrus; BG: basal ganglia; Tha: thalamus; BA: Brodmann area; IFJ: inferior frontal junction; IFS: inferior frontal sulcus; STS: superior temporal sulcus; Ca: caudate; GP: globus pallidus; NAC: nucleus accumbens; Pu: putamen; PM: pre-motor; S: sensory; PP: posterior parietal; O: occipital; PF: pre-frontal; tl: tongue and larynx region; c: caudal; v: ventral; d: dorsal; l: lateral; m: medial; a: anterior; p: posterior; op: opercular.

Notably, the primary regions damaged (with n > 4) did not include lateral aspects of the superior temporal gyrus thought to be involved in speech perception and the representation of phonetic features (Mesegarani et al. 2014; Turkeltaub and Coslett 2010) nor did they include middle and inferior temporal regions thought to represent semantic knowledge (Mummery et al. 1999; Visser et al. 2012). According to an embedded processes view of WM (Cowan et al. in press), persisting activation in such regions supports phonological and semantic WM. Regions beyond these would be those that underlie domain-general attentional processes. Thus, to the extent that our findings uncover regions specific to supporting either phonological or semantic WM, they would argue against the embedded processes view.

### Lesion symptom mapping

Regarding the LSM, both the phonological and semantic WM models were statistically significant (phonological WM: cost = 10^8, gamma = 10^-8, *p* = 0.03; semantic WM: cost = 0.1, gamma = 0.001, *p* = 0.02). Figure 3 and Table 1 display the brain regions whose damage (region size > 100mm^3^) significantly related to either phonological (Figure 3B) or semantic WM (Figure 3C). Because the models controlled for the other WM measure, brain regions significantly related to phonological and semantic WM performance were nonoverlapping.

**Table 1.**
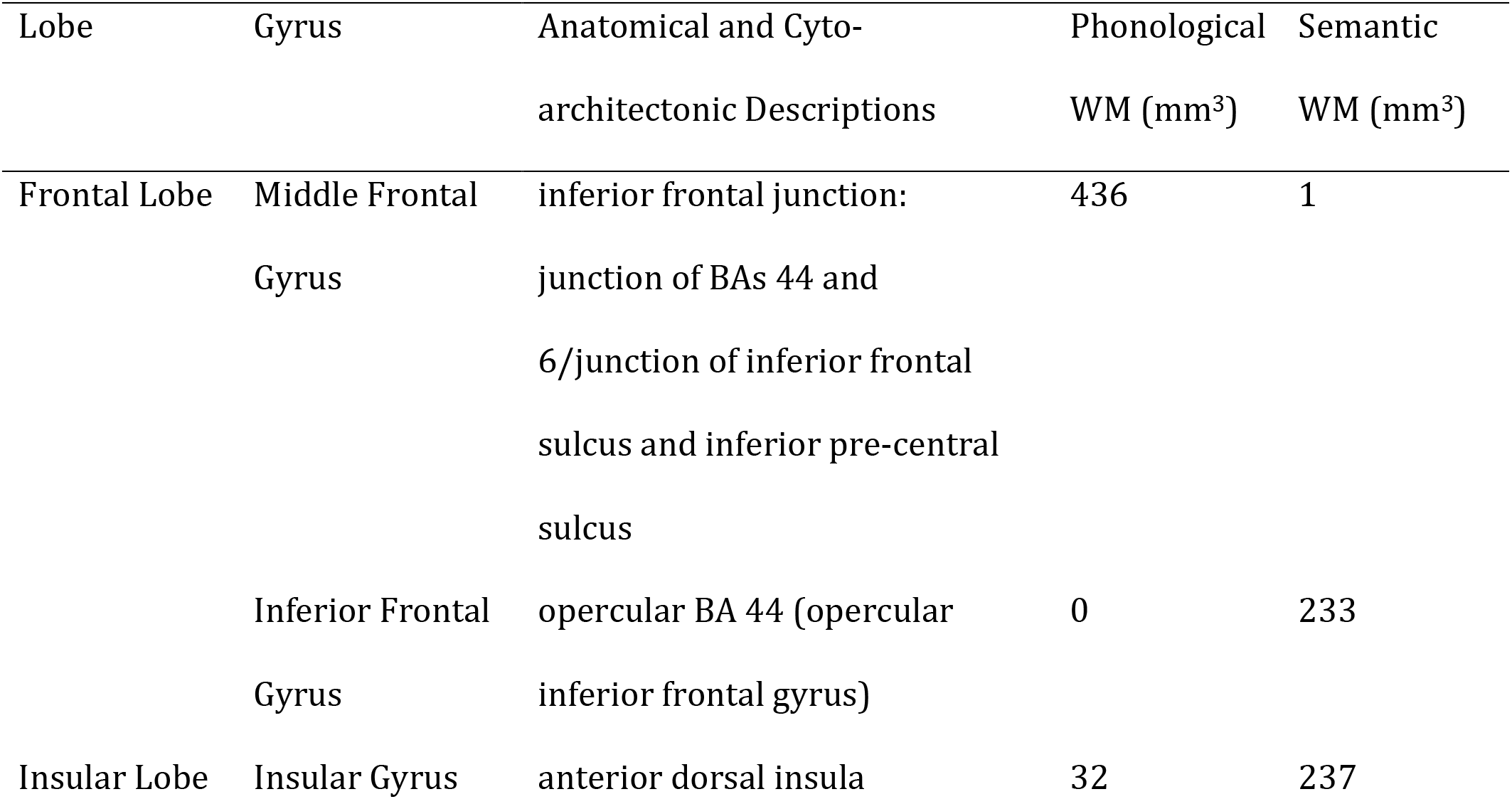
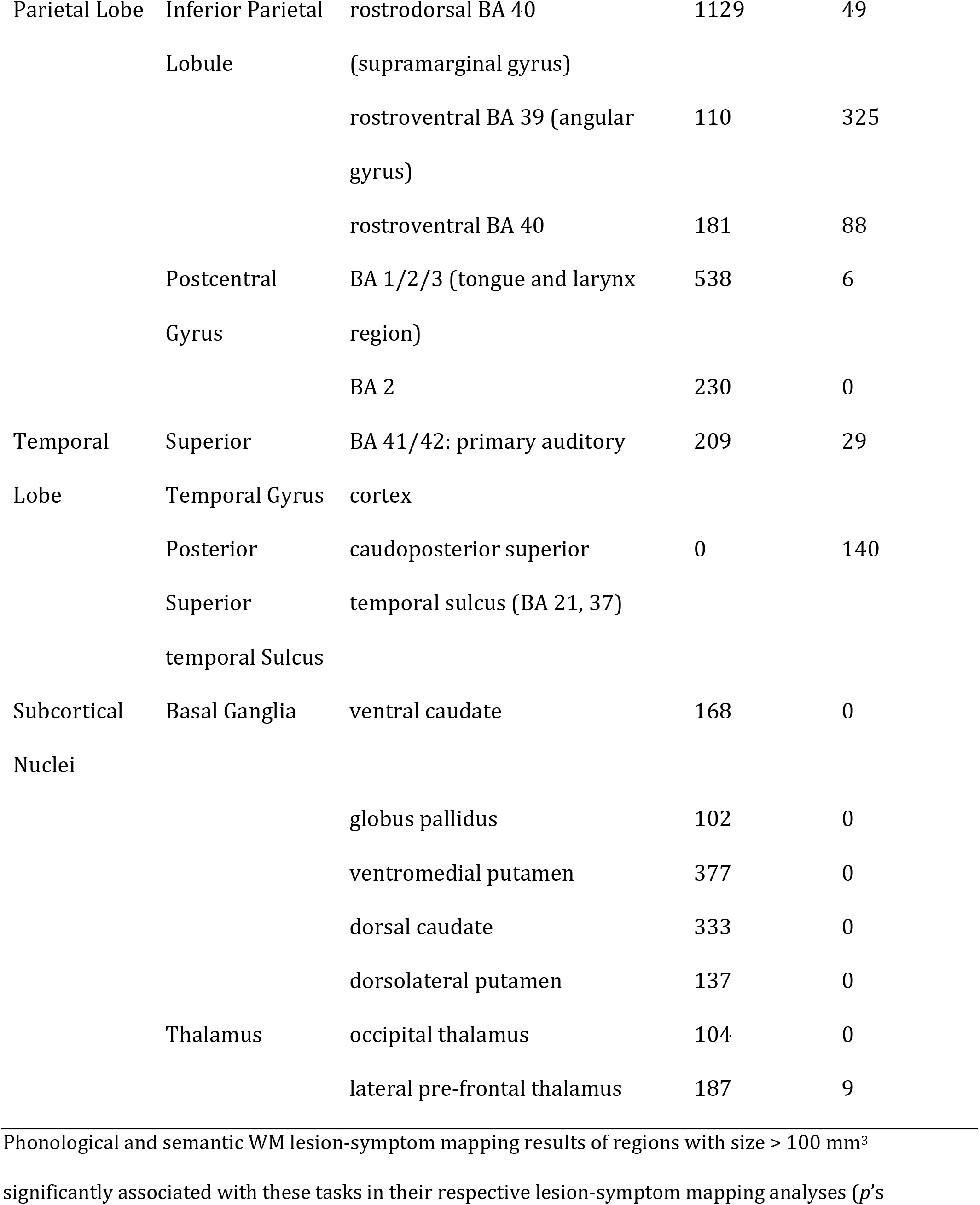
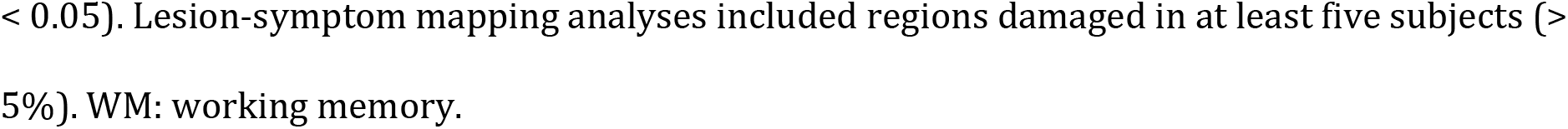
Lesion-Symptom Mapping Results

With respect to phonological WM, the region of the largest size and with the greatest difference between phonological and semantic WM was in the rostrodorsal BA 40 (with suggestion of extension rostroventrally, i.e., SMG; see Table 1), in line with prior findings (Shallice and Vallar 1990; Paulesu et al. 1993). Interestingly, while a region in the superior temporal gyrus related to phonological WM, the region was not in lateral aspects of the STG (BA 21, 22) related to speech perception, but instead in primary auditory cortex (BA 41,42). The other cortical regions specific to phonological WM are all plausibly related to rehearsal, including the inferior frontal junction (junction of BAs 44 and 6, a region joining the inferior frontal sulcus (between the IFG and MFG) and the inferior precentral sulcus), and the post-central gyrus (BAs 1, 2, 3), which includes tongue and larynx regions. A number of areas (basal ganglia and thalamus) were also found, all of which have been implicated in articulatory aspects of speech production (e.g., Bohland and Guenther 2006).

With respect to semantic WM, a region in a somewhat more anterior part of the left IFG (opercular region of BA44) was obtained relative to that for phonological WM. A temporal lobe region bordering the posterior superior temporal sulcus (BA 21,37) was found, which is often implicated in the mapping from phonology to semantics (Okada and Hickok 2006; Wilson et al. 2018). As predicted, an inferior parietal region was obtained in BA39 (AG) which was thus differentiated from the inferior parietal SMG region obtained for phonological WM. The only other region related to semantic WM was the anterior dorsal aspect of the insula. While there is controversy regarding the functional roles of different subregions of the insula, some evidence suggests that anterior dorsal regions are involved in more cognitive processes, including semantic processing (Deen et al. 2010; Ardila et al. 2014)

## Discussion

We examined whether damage to different brain regions caused phonological and semantic WM deficits in a large group of participants during the acute stage of stroke. We avoided the confound of reorganization of function while also controlling for the degree of phonological input processing and access to semantic knowledge. Multivariate lesion symptom mapping (LSM) revealed distinct regions underlying phonological and semantic WM. Although damage to both frontal and parietal lobules reduced WM capacity, the specific regions necessary for each WM capacity differed and did not include regions supporting phonological or semantic long-term knowledge. These results are consistent with a multicomponent view of WM, where functionally and anatomically distinct buffers maintain phonological and semantic information independent from the long-term memory of domain-specific representations.

### Phonological WM

The region with the largest damage associated with reduced phonological WM in comparison to semantic WM was the supramarginal gyrus (SMG) in the inferior parietal lobe. Based on neuroimaging and neuropsychological findings, this SMG region has often been postulated to support the storage of phonological information (Paulesu et al. 1993; Shallice and Papagno 2019; Yue et al. 2018). Other frontal and subcortical regions related to phonological WM were, for the most part, regions carrying out articulatory and motor planning processes, most likely due to their involvement in subvocal rehearsal. Two frontal regions often ascribed a role in motor planning in functional neuroimaging studies (e.g., Chein and Fiez 2001) were also observed here (supplementary motor, posterior IFG). In addition, several subcortical regions were found, all of which have been implicated in motor planning and motor control (Boland and Guenther 2006; Crosson et al. 2003). A few prior LSM studies have reported the involvement of subcortical regions (caudate in Leff et al. 2009, and caudate and putamen in Ivanova et al. 2018), but attributed their role to supporting executive functions involved in short-term memory performance. While we cannot rule out this possibility, both regions have also been found to be involved in simple tasks with little executive function demand involving the production of syllable sequences (Bohland and Guenther 2006). The current study uncovered a number of other subcortical regions not reported in other studies. It is possible that our research was able to detect the involvement of these regions due to testing at the acute stage, whereas other studies have been carried out at the chronic stage (Baldo et al. 2012; Ghaleh et al. 2020; Ivanova et al. 2018; Leff et al. 2009). There is evidence that individuals with subcortical lesions are likely to recover language functions (Demonet 1997; Heiss et al. 1999), thus, limiting the ability to detect the involvement of these regions in supporting phonological WM in a chronic sample.

Interestingly, reduced phonological WM capacity was also related to damage to sensory regions: 1) a somatosensory region representing the lips and larynx, and 2) primary auditory areas (Heschl’s gyrus; BAs 41,42). If an fMRI study of WM revealed these activations, one might suppose that these were the consequence of implicit (or perhaps explicit) motor execution. Thus, one might hypothesize that these regions might be activated during rehearsal but were not necessary for it. However, in the LSM framework, the association of damage to these regions to reduced WM capacity suggests a necessary role. Such a necessary role might be accommodated on the grounds that a somatosensory target is needed to guide motor movements (or, in this case, covert motor movements involved in rehearsal), a proposal that is consistent with the model of Walker and Hickok (2016), which bridges speech production and motor control models. The assumption of a role of somatosensory targets is common in motor theories in the visual-motor domain, supported by a range of findings from humans and non-human animals, and has more recently been extended to speech production (see Hickok 2012 for a review). In contrast, that damage to primary auditory regions reduced phonological WM capacity was somewhat more unexpected. A possible but unlikely explanation is that damage to such regions impaired speech perception, which played a greater role in phonological than semantic WM. However, it is unclear why that should be the case, since one might have expected the small set of digits to be more easily recognized than the words in the category probe task, which varied from trial to trial. More critically, the multivariate LSM analyses controlled for speech perception abilities, by factoring out performance on phonologically related trials in picture-word matching. Instead, we speculate that the involvement of primary auditory regions might be consistent with theories postulating that a match between articulatory and anticipated auditory targets is also used in modulating motor control (Hickok 2012). Overall, for phonological WM, the set of regions hang together as a network plausibly involved in phonological storage and covert rehearsal processes requiring motor planning and execution, though some questions remain regarding the role of the observed somatosensory and primary auditory regions.

### Semantic WM

For semantic WM, damage to a smaller number of regions was associated with deficits. Previous neuropsychological and neuroimaging findings had suggested a critical role for the left inferior/middle frontal region. Although an inferior frontal and insular region was obtained, it was somewhat more posterior than anticipated based on prior fMRI results of semantic WM (Martin et al. 2003; Hamilton et al. 2009). The region that was uncovered (opercular LIFG, BA44) has been implicated in some studies as being involved in semantic selection (Heim et al. 2009; Martin and Chao 2001) and in other studies as playing a role in a semantic maintenance process termed “refreshing” (Johnson et al. 2005; see subsequent section for further discussion). Lesion coverage was not extensive in more anterior aspects of the left inferior frontal and middle frontal gyri, thus limiting our ability to detect the involvement of more dorsal or anterior regions. In posterior areas, a parietal region in the AG (BA 39) and a pSTS (posterior superior temporal sulcus) region (BA21, 37) were obtained. Necessity of the AG for semantic WM is consistent with considerable neuroimaging evidence of its role in semantic processing and, specifically, in semantic integration – that is, the integration of word meanings (Binder et al. 2009; Humphries et al. 2006; Bemis and Pyllkanen 2013). Semantic integration (e.g., integrating noun-noun, “apple core”, and verb-noun, “throw dart”, combinations) would seem to necessarily draw on semantic WM to maintain the two concepts such that an appropriate integration could be carried out. Thus, we are suggesting that the same capacity that supports the maintenance of semantic representations for unrelated words in word lists is drawn upon to hold the meanings of words in sentences prior to their integration. With respect to the pSTS, damage to a similar pSTS locus after acute stroke in a subset of patients studied here was associated with reduced ability to produce nouns and increasingly complex word combinations during a narrative production task (Ding et al. 2020). In a behavioral analysis of another subset of these patients, semantic WM was necessary for producing increasingly complex word combinations (Martin and Schnur 2019). Elsewhere, a recent fMRI study investigating spoken and written narrative comprehension in non-brain damaged individuals (Wilson et al. 2019) revealed the involvement of specific subregions of the pSTS in phonological, lexical, and semantic processes. Based on their findings, they argued that the dorsal posterior region of the STS represented phonological and orthographic lexical representations, whereas the ventral posterior region supported higher level language processes involved in semantic and syntactic processing. Thus, one might postulate that the pSTS region uncovered in the semantic WM task is required either for linking lexical phonological representations with semantic representations or with maintaining word meanings during phrase integration in comprehension and in holding several word meanings during phrase construction in production^4^.

Thus, in general, our results converged with prior findings implicating the left IFG (e.g., Martin et al. 2003; Lewis-Peacock et al. 2012; Shivde and Thompson-Schill 2004) and the angular gyrus (Humphries et al. 2006; Yue et al. 2018) in semantic WM. While the current findings cannot distinguish the role of these two regions, other findings in the literature suggest that the LIFG region is involved in retrieving or refreshing semantic representations (Heim et al. 2009: Johnson et al. 2005; Martin and Chao 2001), whereas the AG is involved more directly in maintaining semantic representations to support meaning integration (Humphries et al. 2006). The results showing the involvement of the pSTS are more novel but may underlie the maintenance of several word meanings during complex phrase construction (Ding et al. 2020). Further investigation is needed to determine if the engagement of this region replicates and what its distinctive role might be.

### Rehearsal vs. Refreshing

As discussed previously, several regions involved in motor control and articulatory planning were found to support phonological WM performance, likely due to their involvement in inner rehearsal. Given that the LSM analyses controlled for the other WM component, these results imply that articulatory rehearsal processes were more important for performance on the phonological than the semantic WM task. In the behavioral literature on healthy adults, a separate process, termed refreshing, has been argued to keep semantic representations in mind (e.g., Loiza and Camos 2018; Nishiyama 2018). Refreshing is held to be a process by which recently activated representations are “rethought” and thereby given a boost of activation, with evidence suggesting independence of refreshing from both articulatory rehearsal and retrieval from long-term memory (Loiza and Camos 2018). Neuroimaging studies have suggested that left middle frontal regions are involved in refreshing semantic representations for words, though, in some experiments, BA44 and the insula were also involved (Johnson et al. 2005), coinciding with the left frontal regions found here. Thus, our findings are consistent with claims of different maintenance mechanisms for phonological and semantic WM.

### Relation to Buffer vs. Embedded Process Theories of WM

As discussed in the introduction, some have argued that there are no dedicated regions for maintaining different types of representations in WM. According to the embedded processes view, persisting activations in regions devoted to long-term memory representations in that domain are thought to underlie WM (Cowan et al. 2020). Our findings for both phonological and semantic WM argue against this view. In the phonological domain, the embedded processes view would lead one to predict that lateral aspects of the STG (BA22), which have been found to underlie speech perception and the representation of sublexical phonological codes, would be critical to phonological WM. However, in our sample there were very few individuals with damage to such regions. Thus, while the present results cannot refute the possibility that lateral STG is a necessary component of the network involved in phonological WM, it is clear that it is not sufficient, as many of our patients had substantially impaired phonological WM capacity resulting from damage to regions elsewhere. In particular, a large region in the inferior parietal lobe in the SMG was found to be related to phonological WM, consistent with prior studies implicating this region as a phonological store (e.g., Deschamps et al. 2014; Yue et al. 2018). Most other areas obtained were plausibly related to covert rehearsal (Boland and Guenther 2006; Chein and Fiez 2001), a process specific to maintaining phonological information in WM (Baddeley et al. in press; Loiza and Camos 2018). Although two sensory areas were obtained, these regions might also be engaged as part of the motor rehearsal process in providing sensory targets used to assess motor accuracy (Guenther et al. 2006; Hickock 2012; Walker and Hickok 2016).

In the semantic domain, we also observed varying degrees of impairment in semantic WM, even though our patients did not have damage to middle or inferior temporal regions thought to house long-term semantic representations for objects or to provide a semantic hub for linking together different aspects of concepts (Mesulam et al. 2015; Visser et al. 2012). Instead, the regions uncovered included frontal (BA44) and parietal regions (BA39), which are plausibly involved in retrieving, maintaining, and integrating semantic information (e.g., Helm et al. 2009; Humphries et al. 2006).

## Conclusions

In summary, this study uncovered distinct regions involved in phonological and semantic WM, while controlling for phonological and semantic knowledge. Most regions that were identified in both domains were separate from regions postulated to be involved in regions representing long-term knowledge of phonology or semantics. Moreover, these results were obtained in a large sample of individuals at the acute stage of stroke, thus ruling out confounds due to reorganization of function. The organization of the regions involved in phonological WM seems fairly clear, consisting of regions involved in phonological storage and motor processes involved in rehearsal (including regions representing sensori-motor targets for articulation). This is the first LSM study of semantic WM, and while distinct frontal and parietal regions were uncovered, future work will be needed to elucidate how these regions interact in supporting semantic WM.

## Funding

This work was supported by an award to the Baylor College of Medicine from the National Institute on Deafness and Other Communication Disorders (R01DC014976).

## Acknowledgements

The authors wish to thank Chia-Ming Lei, Danielle Rossi, and Miranda Brenneman for data collection. We gratefully acknowledge and thank our research subjects and their caregivers for their willingness to participate in this research. This work was presented at the Academy of Aphasia (2020).

## Footnotes

1 Traditional views of memory associated phonological representations with short-term memory and semantic information with long-term memory (e.g., Baddeley 1966). However, many findings indicate an influence of long-term representations of phonology on immediate recall of words and nonwords – for example, better recall when the items are phonologically similar to more words in the language and a separate influence of individual phoneme frequency (e.g., Thorn & Frankish 2005). Thus, as in Figure 1, both phonological and semantic WM can be thought of involving the use of long-term knowledge over the short-term.

2 These studies showed a preservation of gist rather than exact wording in sentence repetition for patients with a phonological WM deficit (e.g., Martin et al. 1994) or an auditory-verbal short-term memory deficit (Saffran & Marin 1975). Although the terminology for their deficits was different, it seems highly likely that both patients had similar WM deficits, as they overlapped on several features, including fluent speech and better WM performance with visual than auditory presentation for word list recall. The preservation of gist implies that despite their WM deficits, they could rapidly derive the meaning of sentences and use that meaning to reconstruct the sentence when the phonological code was no longer available to support recall.

3 The embedded processes account shares some similarities to the earlier levels of processing approach of Craik and Lockhart (1972) in that both emphasize processes rather than storage buffers for WM for maintaining information in mind. However, the embedded processes account does not ascribe to notions of depth of processing determining long-term recall, instead attributing findings of better long-term recall with semantic processing to the relative degree of interference among semantic vs. among phonological representations (Cowan et al. 2020). Of more relevance to the issues described here, both views would lead to the prediction that the same brain regions involved in long-term memory representations and processing in a given domain should support WM in that domain.

4 Lambon Ralph and colleagues (2017) have argued for a semantic control system encompassing frontal and parietal regions that is engaged in accessing and manipulating semantic representations. Thus, one might question the extent to which the regions we uncovered map onto the regions they suggest support semantic control. According to a meta-analysis of neuroimaging studies on semantic control (Noonan et al. 2013), the region closest to our pSTS region was a posterior middle temporal gyrus region; however, the peak activation point was more ventral and posterior than our pSTS region, with no apparent overlap of their cluster and our region. Similarly, with regard to our regions in the AG, their AG region was much more medial and superior to our AG regions. In frontal regions, they uncovered a very large region encompassing most middle and inferior frontal gyrus regions, and thus it would be difficult for our region not to overlap theirs. We would also note, however, that behavioral work has called into question the basis for claims about a distinct semantic control system (Allen, Martin, & Martin, 2012; Chapman, Hasan, Schulz & Martin, 2020).

## Notes

### Competing Interest Statement

The authors have declared no competing interest.

### Summary of Updates

Figure 1 was added to explain the working memory model. The method of category probe and digit span scoring was specified. The method of lesion-symptom mapping was specified. Figure 4 was revised to highlight the working memory related regions. Four footnotes were added to explain some important questions.

